# A high-throughput screening approach to discover potential colorectal cancer chemotherapeutics: Repurposing drugs to disrupt 14-3-3 protein-BAD interactions

**DOI:** 10.1101/2023.12.14.571727

**Authors:** Siyi He, Luis Delgadillo Silva, Guy A. Rutter, Gareth E. Lim

## Abstract

Inducing apoptosis in different types of cancer cells is an effective therapeutic strategy. However, the success of existing chemotherapeutics can be compromised by tumor cell resistance and systemic off-target effects. Therefore, the discovery of pro-apoptotic compounds with minimal systemic side-effects is crucial. 14-3-3 proteins are molecular scaffolds that serve as important regulators of cell survival. Our previous study demonstrated that 14-3-3ζ can sequester BAD, a pro-apoptotic member of the BCL-2 protein family, in the cytoplasm and prevent its translocation to mitochondria to inhibit the induction of apoptosis. Despite being a critical mechanism of cell survival, it is unclear whether disrupting 14-3-3 protein:BAD interactions could be harnessed as a chemotherapeutic approach. Herein, we established a BRET-based high-throughput drug screening approach (Z’-score= 0.52) capable of identifying molecules that can disrupt 14-3-3ζBAD interactions. An FDA-approved drug library containing 1971 compounds was used for screening, and the capacity of identified hits to induce cell death was examined in NIH3T3-fibroblasts and colorectal cancer cell lines, HT-29 and Caco-2. Our *in vitro* results suggest that terfenadine, penfluridol, and lomitapide could be potentially repurposed for treating colorectal cancer. Moreover, our screening method demonstrates the feasibility of identifying pro-apoptotic agents that can be applied towards conditions where aberrant cell growth or function are key determinants of disease pathogenesis.

## Introduction

Apoptosis, or programmed cell death, is a highly regulated process of cell suicide. During apoptosis, cells break down into apoptotic bodies and are eventually engulfed by phagocytes like macrophages and neutrophils^[1]^. With limited leakage of a cell’s content into the extracellular environment, apoptosis can minimize the damage to surrounding cells. The identification of molecules that safely and selectively induce apoptosis holds significant potential in treating a variety of conditions, such as cancer, infectious diseases, and autoimmune disorders^[2–4]^.

Colorectal cancer (CRC) ranks as the second most deadly cancer and accounted for about 9.4% of cancer-induced mortality in 2020^[5]^. CRC carcinogenesis originates from either the colon or the rectum, and most malignant adenomas develop initially from benign polyps^[6]^. Current therapies for CRC involve chemotherapy, radiation therapy, and surgery; however, resistance to existing chemotherapeutics often leads to poor clinical outcomes^[7]^. Accordingly, it remains important to discover novel compounds that induce apoptosis in CRC cells.

An example of an chemotherapeutic that exploits the intrinsic pathway of apoptosis is Venetoclax (ABT-199). ABT-199 is a specific inhibitor of BCL-2, an anti-apoptotic member of the BCL-2 protein family^[8]^. The effectiveness of ABT-199 in treating chronic lymphocytic leukemia (CLL) and acute myeloid leukemia (AML) highlights the potential of targeting the actions of the BCL-2 protein to treat cancers. We previously demonstrated that 14-3-3ζ, a member of the 14-3-3 scaffold protein family, plays an essential role in maintaining the survival of MIN6 insulinoma cells through its inhibitory actions on pro-apoptotic BCL-2 protein, such as BAD and BAX^[9]^. In healthy murine cells, 14-3-3 proteins sequester BAD in the cytoplasm by interacting with phosphorylated Ser112 and Ser136 residues on BAD^[10]^. However, the induction of cell death or prolonged cell stress results in these serine residues becoming dephosphorylated, leading to the dissociation of 14-3-3 protein:BAD complexes. This allows BAD to translocate to the outer mitochondrial membrane, where it interacts with other BCL-2 proteins, such as BCL-2 and BCL-xL, to initiate apoptosis^[11–13]^.

14-3-3 proteins have been found to play important roles in cancer cell survival^[14]^. Alterations in 14-3-3 protein expression, especially the 14-3-3ζ isoform, have been observed in a variety of cancers, such as those of colon, breast, lung, and pancreas^[15]^, and overexpression of 14-3-3ζ may mediate tumor resistance to chemotherapy due to its anti-apoptotic functions^[16]^. Depletion of 14-3-3ζ has been found to induce the apoptosis of CRC cells *in vitro* and *in vivo*^[17]^, and suggesting that identifying or developing novel disruptors of 14-3-3 protein:BAD protein-protein interactions (PPIs) might represent a promising approach towards the treatment of various cancers, including CRC.

Drug development involving *de novo* synthesis and validation of novel chemical entitites is an incredibly expensive and time-consuming endeavor that typically yields a low success rate^[18–20]^. From preclinical studies to clinical trials, and ultimately to approval by the US Food and Drug Administration (FDA), the estimated average cost per drug was over $1.5 billion between 2009 to 2018, with a development time of up to over 20 years^[18, 19]^. However, even with such huge investments, only 10% of drugs that enter phase I clinical trials are approved^[20]^. Given the challenges of new drug development, repurposing already approved compounds for new indications could be an attractive strategy, as it saves both time and costs^[21–23]^. To date, the most comprehensive compound screen aimed at identifying disruptors of 14-3-3 protein:BAD PPIs was performed with a time-resolved fluorescence resonance energy transfer (TR-FRET)-based approach, and while 16 hits were discovered from over 52100 examined compounds, an important caveat was that this assay was based on a cell-free system involving recombinant proteins^[24]^.Thus, it was not possible to discern if identified hits would act via receptor-mediated pathways or be transported into a cell to disrupt 14-3-3 protein:BAD PPIs.

Herein, we have developed an innovative BRET(bioluminescence resonance energy transfer)-based biosensor to detect 14-3-3 protein:BAD PPIs in intact, living cells^[25]^. Using this sensor and an FDA-approved drug library containing 1971 compounds, we first identified 101 hits through a high throughput screening (HTS) approach in NIH-3T3 fibroblasts. We next evaluated the capacity of these hits to induce cell death, and 41 compounds emerged as potential candidates. Based on their original indications and routes of administration, we further selected 13 drugs to assess their effectiveness in inducing apoptotic cell death in CRC cell lines, HT-29 and Caco-2. Our screening workflow has identified terfenadine, penfluridol, and lomitapide as potent candidate molecules that can potentially be repurposed as chemotherapies towards inducing CRC cell death.

## Materials and Methods

### BRET sensor construction

The original plasmids containing 14-3-3ζ, BAD, and BAD mutants were kind gifts from Dr. Herman Spaink and Dr. Aviva M Tolkovsky, respectively^[26, 27]^. To conjugate mTurquoise, a cyan fluorescent protein (CFP) to the C- or N-termini of 14-3-3ζ, 14-3-3ζ was subcloned into pmTurquoise2-N1 (Addgene, Massachusetts, USA; plasmid # 54843) using restriction enzymes EcoRI (NEB, Massachusetts, USA; # R0101S) and AgeI (NEB; # R0552S) and pmTurquoise2-C1 (Addgene; plasmid # 60560) using EcoRI and KpnI-HF (NEB; # R0145S), respectively. pmTurquoise2-N1 and pmTurquoise2-C1 were gifts from Michael Davidson and Dorus Gadella^[28]^. For the conjugation of Renilla luciferase-8 (Rluc8) to the C-termini of 14-3-3ζ, Rluc8 was subcloned from pcDNA-Rluc8 (a kind gift from Dr. Jace Jones-Tabah and Dr. Terry Hébert, McGill University) and used to replace the mTurquoise of constructed 14-3-3ζ-mTurquoise using AgeI and NotI-HF (NEB; # R3189S). To conjugate Rluc8 to the N-termini of 14-3-3ζ, Rluc8 was subcloned to the original 14-3-3ζ-containing plasmid using EcoRI and KpnI-HF^[26]^. Specific primers were used to generate truncated forms of BAD, which were then subcloned into pmCitrine-C1 (Addgene; plasmid #54587) and pmCitrine-N1 (Addgene; plasmid #54594) using EcoRI and BamHI (NEB; #R0136S). This process attached mCitrine, a yellow fluorescent protein (YFP), to the N-termini and C-termini of BAD, respectively. pmCitrine-C1 and pmCitrine-N1 were gifts from Robert Campbell, Michael Davidson, Oliver Griesbeck, and Roger Tsien^[29]^. To construct bi-directional plasmids, 14-3-3ζ with different conjugated molecules and BAD variants conjugated with mCitrine were subcloned to the multiple cloning site (MCS-2) using EcoRI and XbaI (NEB; # R0145S) and the MCS-1 using MluI-HF (NEB; # R3198S) and SalI-HF (NEB; # R3138S) of pBI-CMV1 vector (Takara, Shiga, Japan; # 631630), respectively. Primers were designed with SnapGene Viewer (7.1.0) and synthesized by Integrated DNA Technologies (IDT, California, USA). Phusion® High-Fidelity DNA Polymerase (NEB; # M0530S) was used for PCR amplification. QIAquick Gel Extraction Kit (Qiagen, Hilden, Germany; # 28704) was used to recover DNA products from agarose gels. T4 DNA ligase (NEB; # M0202S) was used to insert genes into vectors. All restriction enzymes were purchased from NEB. Primer sequences are listed in Table 1.

**Table 1.**
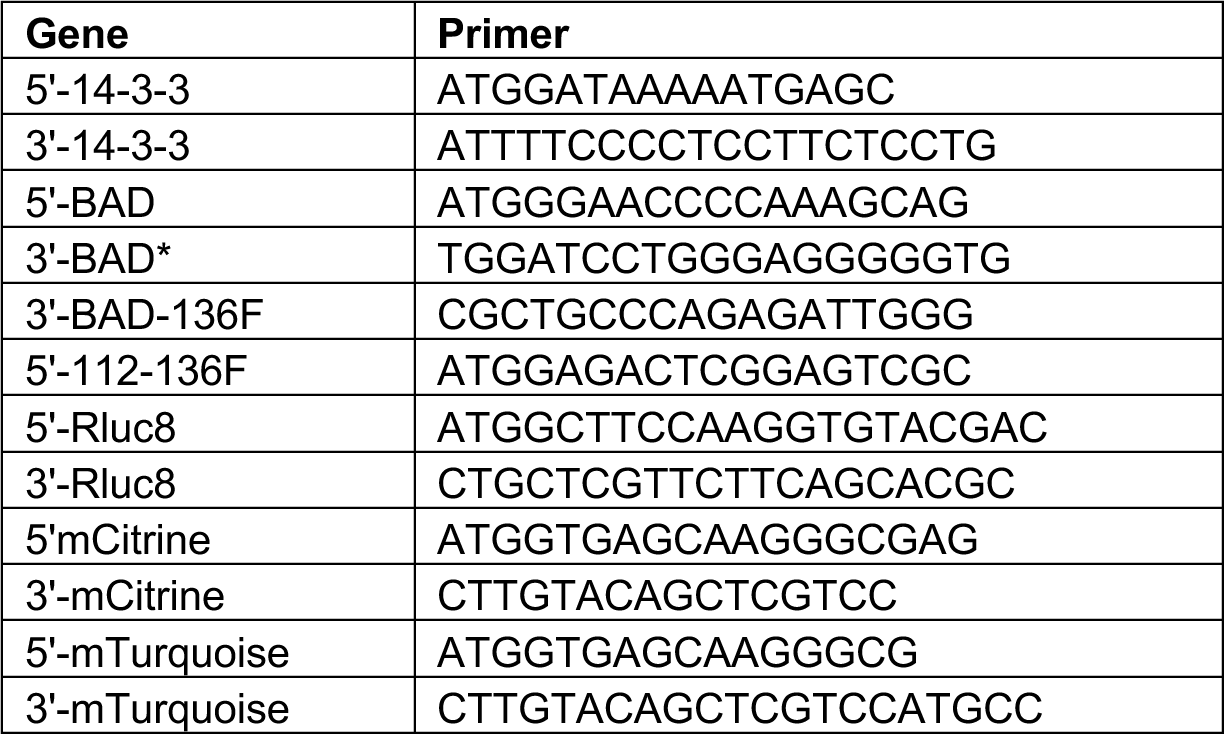
Primers and restriction enzymes for the construction of BRET sensor.

### Cell Culture

NIH-3T3 cells were kindly provided by Dr. Marc Prentki (CRCHUM, Montreal, Canada) and maintained in 25 mM glucose DMEM (Gibco, Massachusetts, USA; # 11995065) supplemented with 10%FBS (Gibco; # 12483020) and 1% penicillin-streptomycin (Gibco; # 15140122). HT-29 and Caco-2 cells were kind gifts from Dr. Petronela Ancuta (CRCHUM, Montreal, Canada) and maintained in Advanced MEM (Gibco; # 12492013) supplemented with 20% FBS and 1% penicillin-streptomycin or McCoy 5A media (Gibco; # 16600082) supplemented with 10%FBS and 1% penicillin-streptomycin, respectively. All cells were cultured in a humidified incubator at 37°C with 5% CO_2_ and passaged upon reaching 70%-80% confluency. All studies were repeated with cells at different passages to ensure reproducibility.

### Plasmid transfection

Plasmid DNA was amplified in Subcloning Efficiency DH5α competent cells (Invitrogen, Massachusetts, USA; # 18265-017) and extracted using QIAprep Spin Miniprep Kit (Qiagen; # 27104) or PureLink™ HiPure Plasmid Maxiprep Kit (Invitrogen; # K210007), following the manufacturer’s protocol. The purity and concentration of each plasmid were assessed with a NanoDrop™ Lite Spectrophotometer (Invitrogen). Two days before transfection, cells were plated in 12-well plates with a density of 30,000 cells/well or proportionally adjusted based on the surface area of different well sizes. Plasmids were transfected with Lipofectamine™ 3000 Transfection Reagent (Invitrogen; # L3000015), according to the manufacturer’s instructions.

### Confocal imaging

Confocal images were acquired with a Leica TCS-SP5 inverted microscope using an HCX PL APO CS 100x/1.4 Oil objective with the Las-AF software. Excitation was performed using the 458nm and 514nm laser lines of an Argon laser for CFP and YFP, respectively, and a 633nm HeNe laser for TOPRO-3. Detection bandwidth were set to 468-500nm for CFP using a HyD detector under the Standard mode, 524-623nm for YFP using a PMT and 643-748nm for TOPRO-3 using a HyD detector under the Standard mode. A sequential acquisition consisting of CFP and YFP in sequence 1, and TOPRO-3 in sequence 2 was performed. A line average 4 was applied for each sequence. Images were acquired at zoom 2.5 with a 400Hz scan speed and final images are 12bits, 1024×1024pixels (axial pixel size of 60nm).

To visualize BAD translocation, 35mm glass bottom dishes (ibidi, Gräfelfing, Germany; # 81158) were coated with 0.1 mg/mL Poly-D-lysine hydrobromide for 2 hours at 37°C. After coating, the dishes were washed twice with sterilized water. Subsequently, cells transfected with BAD-mCitrine were seeded at a density of 20,000 cells per dish and incubated for 24 hours to allow for cell attachment. For live-cell imaging (Leica TCS SP5, Leica Microsystems, Wetzlar, Germany), cells were incubated for 1 hour with the mitochondrial membrane potential dye tetramethylrhodamine ethyl ester (TMRE) (Invitrogen; # T668) at a final concentration of 100nM. Super-resolution images were acquired on a Zeiss (Oberkochen, Germany) LSM-900 Airyscan inverted confocal microscope equipped with an environmental incubation system, laser lines including 488 and 561 nm, and an oil immersion 60×/1.40 Plan achromat objective. The signal from YFP fluorophore (ex. 488 nm, det. 525-542 nm) and TMRE (ex. 561 nm, det. 600-700 nm) were collected in super-resolution mode. Cell viability dye DRAQ (BioLegend, California, USA; # 424101) was added to the medium at room temperature for 30 minutes prior to live-cell imaging.

### High-throughput screening (HTS) of drugs

The Z’-score, which evaluates the robustness of an HTS assay readout, was calculated according to prior research^[30]^. For HTS, 384-well plates were coated with 100ug/mL poly-L-ornithine (Sigma, Missouri, United States; # P3655) overnight at 37°C, followed by rinsing with sterilized water. NIH-3T3 cells were transfected with the BRET sensor two days prior in 10cm dishes and replated into pre-coated 384-well plates at a density of 20 000 cells/well. After 24 hours, the growth media was replaced by Krebs buffer (pH 7.4). A robotic liquid handling system (Eppendorf, Hamburg, Germany; epMotion 5075) was used for the dilution of the drug library (APExBIO, Texas, USA; # L1021) and the addition of drugs to corresponding wells. Each compound was tested at concentrations of 200uM, 20uM, 2uM, and 200nM, whereby cells were exposed to drugs for 3 hours. Coelenterazine-h (NanoLight Technology, Arizona, USA; # 301), the substrate of Rluc8, was added to each well with a final concentration of 5uM. After incubation for 10 minutes, emission from Rluc8 at 460nm and mCitrine at 535nm were measured (Perkin-Elmer, Massachusetts, USA; 1420 Multilabel counter). BRET ratios were calculated by the ratio of emission readings at 535/460 nm^[31]^, and BRET reduction was quantified by normalizing the BRET ratio of drug-treated cells [(BRET (drug)] to the BRET ratio of solvent-treated control cells [BRET (control)], using the formula: [BRET (control) – BRET (drug)] / BRET (control).

### Cell death and apoptosis assays

Cells were plated with a density of 20,000 cells/well into 96-well plates (PerkinElmer; # 6055302) and incubated overnight before the addition of drugs. One hour prior to imaging, 50ng/ml Hoechst 33342 (Invitrogen, # H3570), and 0.5ug/ml Propidium Iodide (Invitrogen; # P3566) were added to wells. To measure apoptosis, AlexaFluor 488-conjugated Annexin V (ThermoFisher, Massachusetts, USA; # A13201) was also added at a dilution of 1:400. A high-content imaging system (PerkinElmer; Operetta CLS), was used to quantify cell death and apoptosis by calculating the number of propidium iodide-positive cells relative to Hoechst-positive cells, and the number of Annexin V-positive cells relative to Hoechst-positive cells, respectively. To assess caspase-dependent apoptosis, cells were pre-treated for 30 minutes with the caspase inhibitor Z-VAD-FMK (BD biosciences, New Jersey, USA; #550377; 20uM) or the negative control Z-FA-FMK (BD biosciences; #550411; 20uM) prior to the incubation of drugs.

### Statistical analysis

R programming (4.3.2) was used for generating scatter density plots. Harmony® high-content analysis software (4.9) was used for processing data collected by the Operetta CLS system. Data were analyzed by GraphPad Prism (version 9.5.0), and the statistical significance of the results was examined by one-way ANOVA with a Dunnett post hoc test or two-way ANOVA with a Tukey post hoc test. Statistical significance in the figures is shown as follows: \**P* < 0.05; \*\**P* < 0.01; \*\*\**P* < 0.001; \*\*\*\**P <* 0.0001. Data are presented as mean ± SEM.

## Results

### Inhibition of 14-3-3 proteins promotes BAD translocation

The canonical mechanism by which 14-3-3 proteins regulate cell survival is through the sequestration of pro-apoptotic BCL-2 proteins, such as BAD, in the cytoplasm^[32]^. To visualize how 14-3-3 protein inhibition impacts BAD localization and subsequent translocation to mitochondria, BAD-mCitrine was transiently expressed in NIH-3T3 cells, followed by incubation with TMRE (tetramethylrhodamine, ethyl ester) to label mitochondria. R18 and FTY720, which are established 14-3-3 protein inhibitors that act through distinct mechanisms of action^[33, 34]^, were used to inhibit 14-3-3 proteins. BAD-mCitrine-expressing cells or control mCitrine cells were exposed to R18 (10 µM) and FTY720 (2 µM) for 24 hours, and localization of mCitrine was visualized by confocal microscopy. A BAD mutant containing S112A and S136A double mutations (BAD-AA), which prevents 14-3-3 protein:BAD PPIs, was used as a positive control. In the absence of 14-3-3 protein inhibitors, BAD-mCitrine was distributed diffusely throughout the cytoplasm (**Figure 1**), and inhibition of 14-3-3 proteins with FTY720 and R18 promoted BAD translocation to TMRE-labelled mitochondria. This redistribution was also observed in cells expressing the BAD-AA-mCitrine mutant. These observations indicate the essential role of 14-3-3 proteins in the cytoplasmic sequestration of BAD.

**Figure 1.**
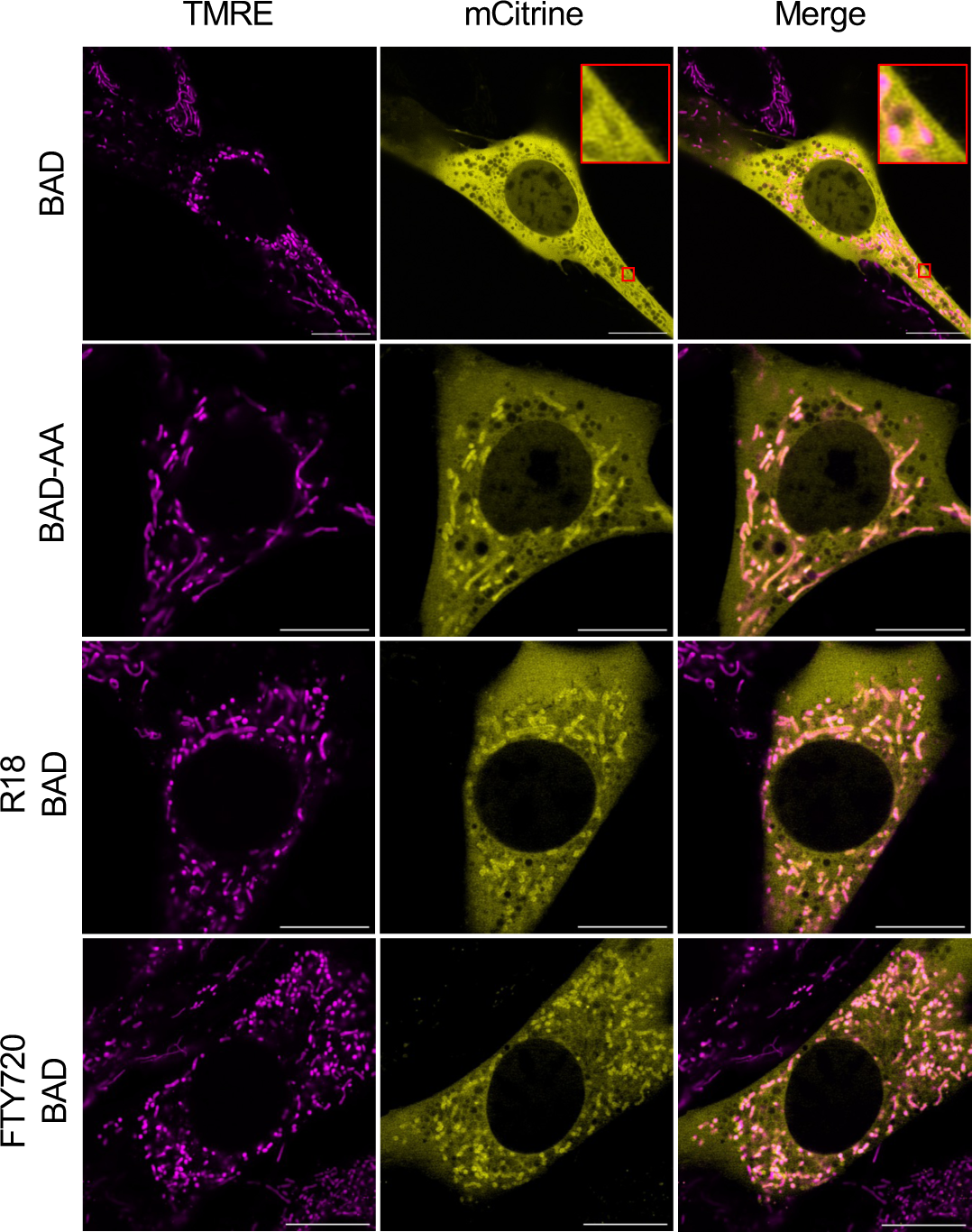
Disrupted 14-3-3 functions promotes BAD translocation. NIH-3T3 cells were made to express mCitrine-conjugated BAD (BAD) or BAD mutants harboring S112A and S136A double mutations (BAD-AA). BAD-expressing cells were treated with either R18 (10uM) or FTY720 (2uM) for 24 hours. The mitochondrial membrane potential sensor TMRE (100 nM) was added to the medium 1 hour prior to live-cell imaging. Representative images were selected from three independent experiments, with at least five images were captured per dish in each experiment. (Scale bar= 10µm)

### Development of a BRET-based living-cell sensor to detect interactions between 14-3-3ζ and BAD

To measure 14-3-3 protein:BAD PPIs in living cells, we focused our efforts on developing a BRET-based reporter whereby 14-3-3ζ and BAD would be conjugated to Rluc8 and mCitrine, respectively (**Figure 2A**). As BRET is highly dependent on the close proximity of the donor (Rluc8) and acceptor (mCitrine), we compared BRET efficiency if mCitrine was fused at the N- or C-termini of full-length murine BAD or truncated versions of BAD (**Figure 2B, Supplementary figure 1A)**. Truncated BAD fragments included BAD-136F, which extends from the N-terminus of BAD to A144, and BAD-112-136F which spanned from M104 to A144. Moreover, we also compared BRET efficiency if Rluc8 was conjugated to the N- or C-termini of 14-3-3ζ (**Figure 2C, D**). The pairing of 14-3-3ζ-Rluc8 with BAD-112-136F-(mC)itrine generated the most robust BRET signal compared to other combinations (**Figure 2C, 2D**).

**Figure 2.**
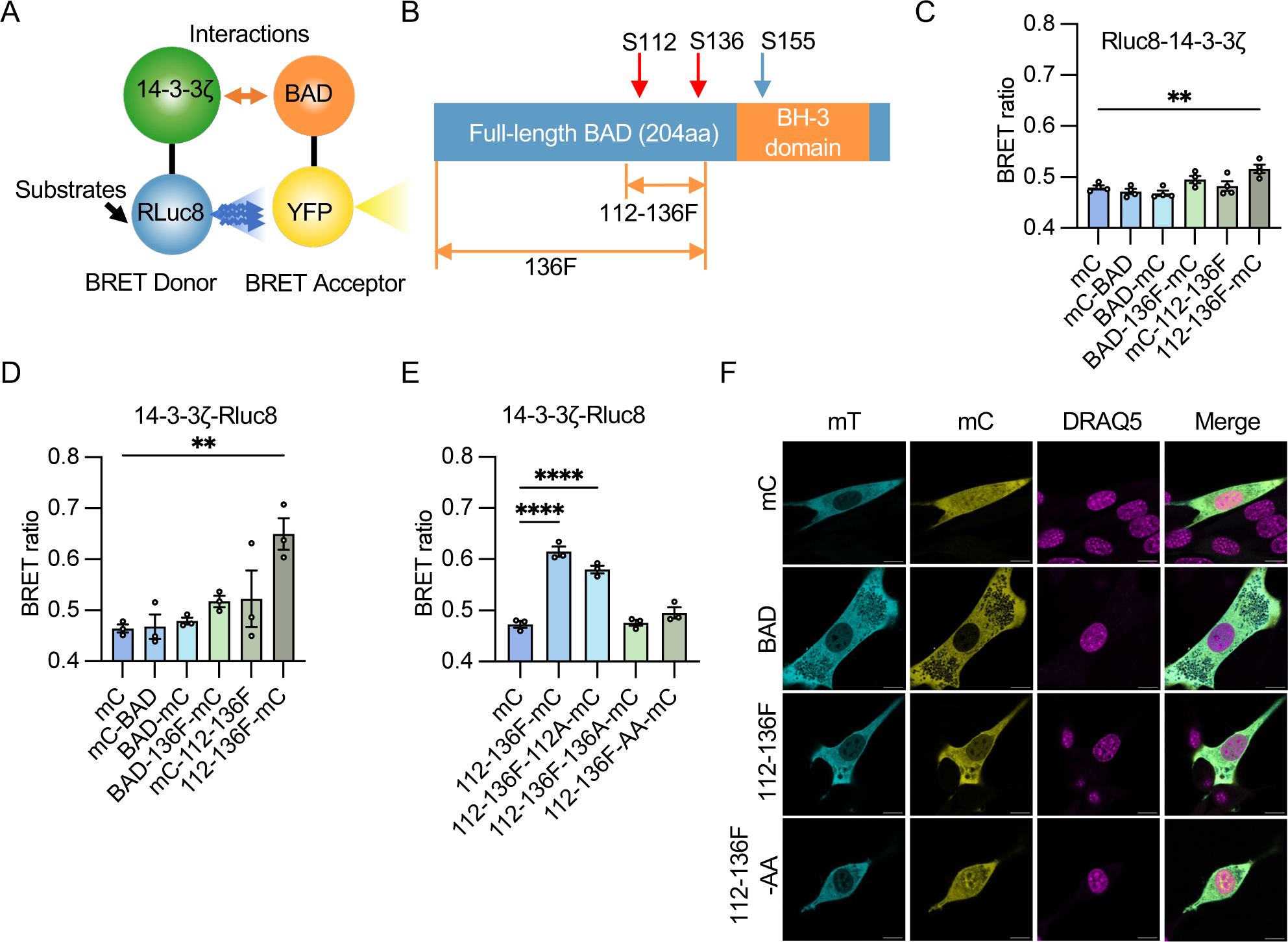
Constriction of a BRET sensor to detect interactions between 14-3-3 and BAD. **(A)** Schematic of the BRET sensor: Rluc8 (BRET donor) and mCitrine (BRET acceptor) are conjugated to 14-3-3ζ and BAD or BAD variants, respectively. The interaction between 14-3-3ζ and BAD or its variants brings the donor and acceptor into close proximity, facilitating BRET. (**B**) Illustration of BAD and truncated forms of BAD. (**C**,**D**) Various pairings of BRET donor and acceptor were assessed. Rluc 8 was conjugated to either the N-termini (Rluc8-14-3-3ζ; C; n=4 per group) or C-termini (14-3-3ζ-Rluc8; D; n=3 per group) of 14-3-3ζ. mCitrine (mC) was linked to the N-termini of BAD (mC-BAD), the truncated BAD spanning Met106 to Ala144 (mC-112-136F), or to the C-termini of BAD (BAD-mC), truncated BAD extending from Met1 to Ala144 (BAD-136F-mC), and the fragment 112-136F (112-136F-mC). (**E**) Single mutation S112A (112-136-F-112A) and S136A (112-136F-136A), and a double mutation combining S112A and S136A (112-136F-AA) were introduced to confirm the interactions between 14-3-3ζ-Rluc8 and 112-136F-mC (n=3 per group). (**F**) In NIH-3T3 cells were co-transfected with 14-3-3ζ-mTurquoise (14-3-3ζ-mT) and either mC, BAD-mC, 112-136F-mC, or 112-136-F-AA-mC, and cell nuclei were visualized with 5 µM DRAQ5. Representative images were selected from three independent experiments, with three images were captured per dish in each experiment. (Scale bar= 10 µm. (Statistical significance was determined using one-way ANOVA with a Dunnett’s post hoc test. *#P* < 0.05 when compared with mC; \**P* < 0.05; \*\**P* < 0.01; \*\*\*\**P <* 0.0001 (C,D,E)).

To confirm the interactions between 14-3-3ζ-Rluc8 and BAD-112-136F-mC, we introduced single serine-to-alanine mutations at either S112 (BAD-112-136F-112A-mC), S136 (BAD112-136F-136A-mC), or a double S112/136AA mutation (BAD-112-136F-AA-mC), as these mutations are known to prevent binding of 14-3-3 proteins to BAD. We found that BAD-112-136F-136A-mC and BAD-112-136F-AA-mC could not elicit a detectable BRET signal, which indicated an inability for 14-3-3ζ-Rluc8 to interact with either BAD variant (**Figure 2E**).

Using confocal microscopy, the subcellular localizations of 14-3-3ζ and BAD-112-136F were determined. Conjugation of (mT)urquoise to 14-3-3ζ revealed that 14-3-3ζ is primarily restricted to the cytoplasm (**Figure 2F**). We next compared the subcellular localization of BAD-mC to BAD-112-136F-mC and found that the fragment was similarly restricted to the cytoplasm. In contrast, BAD-112-136F-AA-mC was distributed throughout the cell, including the nucleus (**Figure 2F**).

With our observations that co-transfection of 14-3-3ζ-Rluc8 and BAD-112-136F-mC plasmids resulted in a detectable BRET signal, we constructed a bi-directional BRET sensor plasmid whereby the donor and the acceptor were expressed at near stoichiometric ratios due to equal promoter activities (**Figure 3A**). Co-expression of BAD-112-136F-mCitrine and 14-3-3ζ-Rluc8 in NIH-3T3 cells resulted in an average of 34.56% increase in BRET compared to the co-expression of 112-136F-AA-mCitrine and 14-3-3ζ-Rluc8 (**Figure 3B**). To test the performance of our bi-directional BRET sensor, we evaluated the capacity of our sensor to detect 14-3-3 protein:BAD interactions with two well-recognized 14-3-3 inhibitors, FTY720 and I,2-5^[35]^. Treatment of sensor-expressing cells with FTY720 and I,2-5 significantly reduced BRET in a dose-dependent manner (**Figure 3C, 3D**). To optimize conditions for the following HTS, we next determined the optimal incubation time and found that 3 hours was required to reduce the magnitude of BRET to the same degree as BAD-112-136-AA, which indicated a maximal effect (**Figures 2E,3E**).

**Figure 3.**
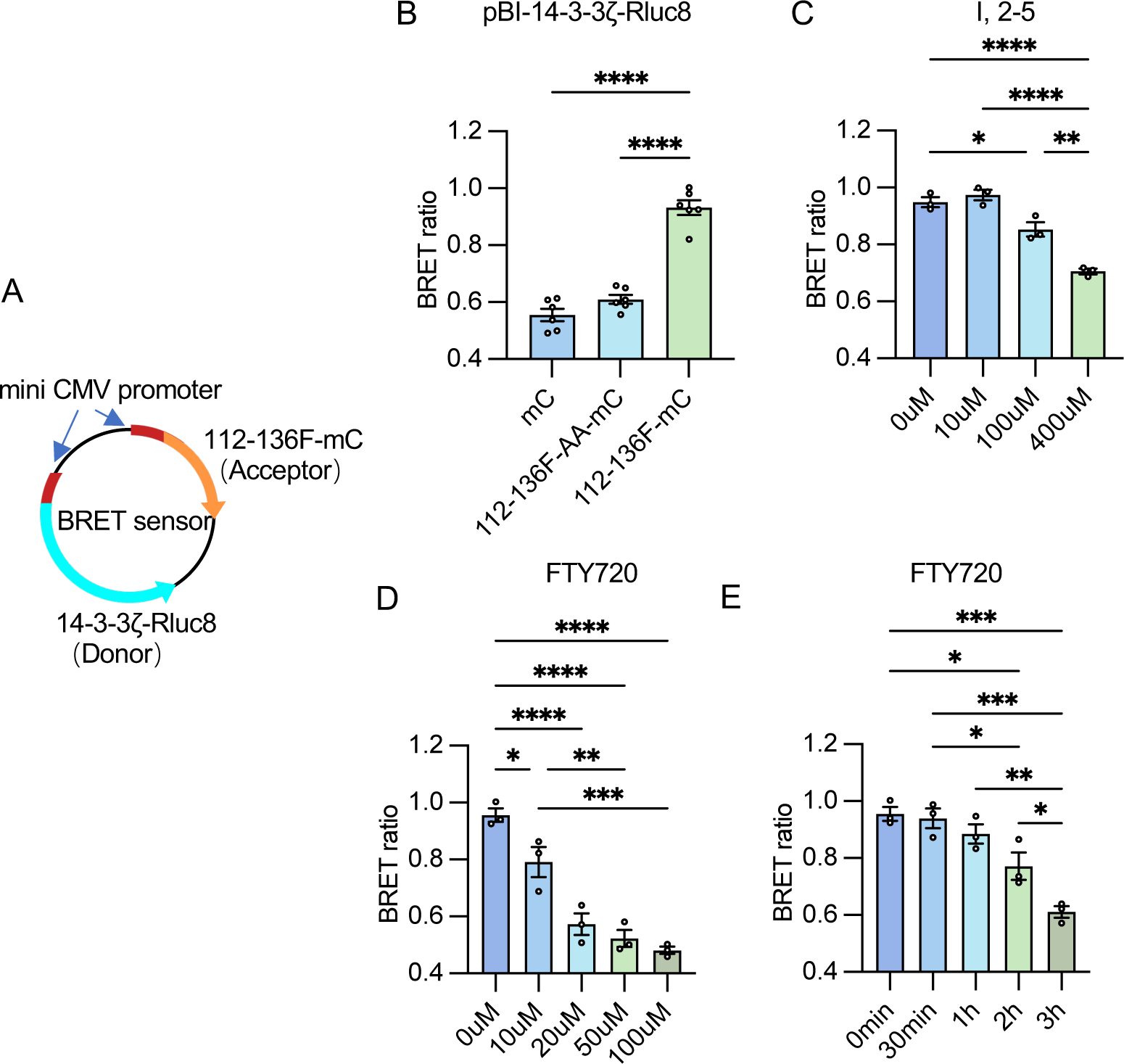
Validation of the BRET sensor. **(A)** Schematic of the BRET sensor design. 14-3-3ζ-Rluc8 and 112-136F-mC were inserted into pBI-CMV1 vector at MCS2 and MCS1, respectively. Plasmids with mCitrine (mC) or 112-136-AA-mC inserted in place of 112-136F-mC were generated as controls(**B**) BRET ratios between 14-3-3(-Rluc8 and mC, 112-136-AA-mC, or 112-136F-mC were compared (n=6 per group). (**C**) A dose response study was conducted with I,2-5 in NIH-3T3 cells expressing the BRET sensor. Measurements were taken 3 hours post-treatment (n=3 per group). (**D**) A dose response study was conducted with FTY720. Data were collected 3 hours post-treatment (n=3 per group). (**E**) A time response study was conducted with FTY720 (20 µM) (n=3 per group). (Statistical significance was determined using one-way ANOVA with a Dunnett’s post hoc test. \**P* < 0.05; \*\**P* < 0.01; \*\*\**P* < 0.001; \*\*\*\**P <* 0.0001)

### Screening for FDA-approved pro-apoptotic drugs

The successful implementation of our BRET-based sensor in living cells permitted the screening of previously-approved drugs (PADs) that could disrupt 14-3-3 protein:BAD interactions (**Figure 4A**). The re-purposing or re-positioning of these drugs could lead to the successful identification of new functions in the induction of cell death. The primary screen was conducted in a 384-well plate format and 1971 compounds were tested (**Figure 4B**). We found the median BRET reduction caused by compounds capable of reducing BRET was approximately 25%, and there were 416, 162, 31, and 16 PADs that reduced the BRET signal by 25% at concentrations of 200 µM, 20 µM, and 2 µM, respectively (**Figure 4B**). We next adapted our screen into 96-well plates to re-screen PADs that were effective at 20 µM, a concentration commonly used in HTS assays^[36]^. Of these, 101 PADs showed consistent results and were further assessed for their capacity to induce cell death in NIH-3T3 cells via Hoechst/propidium iodide incorporation assays (**Figure 4C**). Scatter density plots were used to visualize the relationship between BRET reduction and the induction of cell death at 24 and 48 hours (**Figure 4D, 4E**), and 41 PADs were found to reduce BRET by more than 34%, which is consistant with the reduction in BRET caused by 112-136F-AA-mCitrine (**Figure 3B**), and induce cell death greater than 30% (Zone A; **Figure 4D,E**).

**Figure 4.**
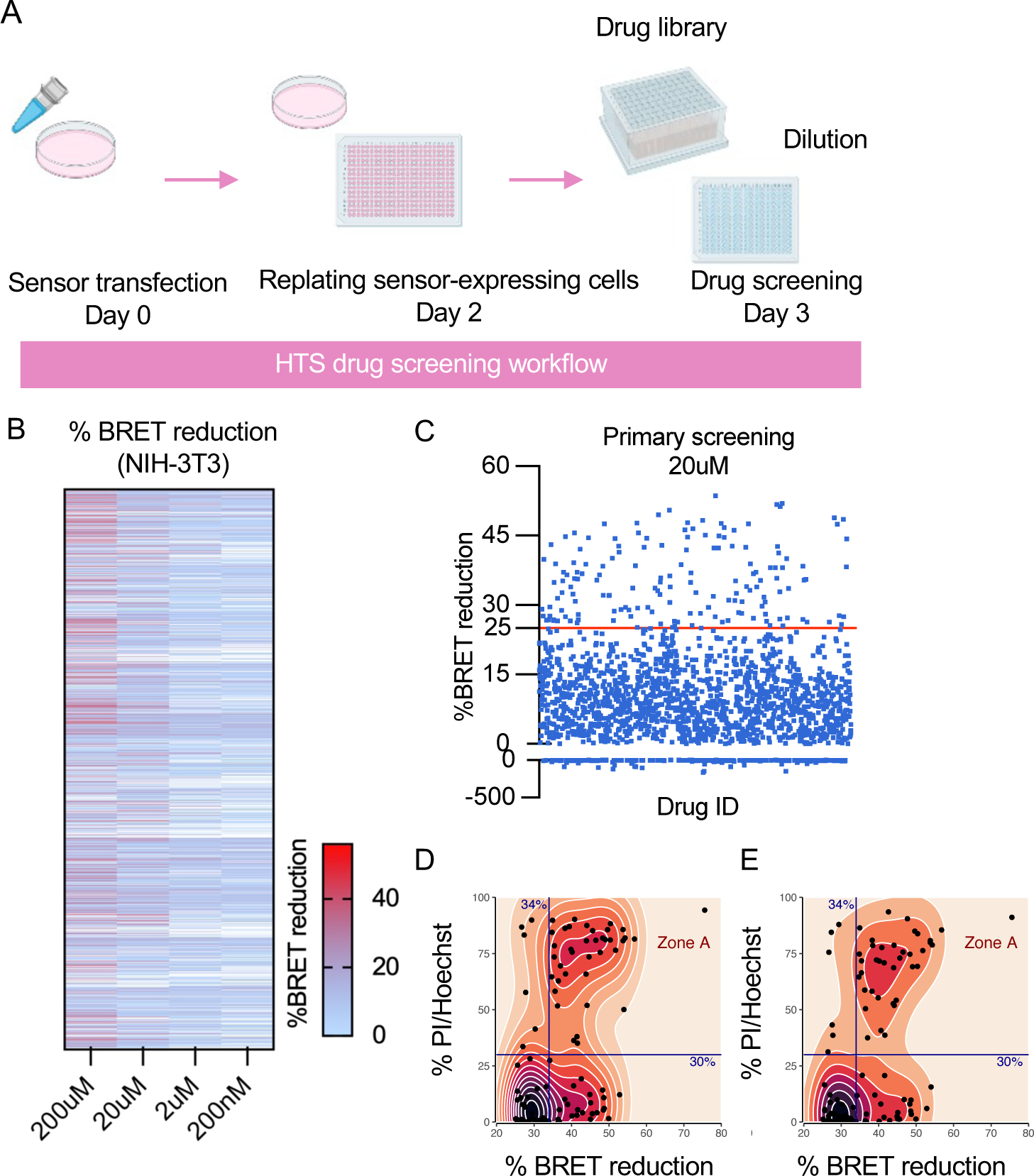
High throughput screening (HTS) for drugs that disrupt 14-3-3:BAD interactions. (**A**)Schematic of HTS workflow. Cells were transfected with BRET sensor or control sensors in 10 cm dishes and allowed 48 hours for sensor expression. Subsequently, cells were harvested and re-plated into 384-well plates at a density of 20,000 cells per well 24 hours prior to drug screening. Diluted previously-approved drugs (PADs) were added to corresponding wells with final concentrations of 200 µM, 20 µM, 2 µM, and 200 nM, respectively. After a 3-hour incubation, the BRET ratio was measured Each PAD was assessed twice at each concentration, but only the highest value for each concentration was recorded. (**B**) The heatmap shows BRET reduction for 1971 PADs at concentrations of 200 µM, 20 µM, 2 µM, and 200nM. (**C**) Subsequent screenings were conducted at 20 µM, And PADs were selected based on the median BRET reduction (26.80%) of those that demonstrated a BRET reduction greater than zero. To include the hits near this threshold, a BRET reduction of 25% was set as the selection criterion for primary screening. This led to the selection of 162 PADs for re-screening in a 96-well plate format. (**D, E**) 101 hits were further selected to assess their capacity to induce cell death in NIH-3T3 cells. Hoechst/propidium iodide incorporation assays were used to examine cell death, calculated as the ratio of propidium iodide-positive cells to Hoechst-positive cells. Cell death data were collected from three independent experiments, each with duplicate measurements. Scatter density plots are used to visualize the relationship between drug-induced BRET reduction and their capacity to induce cell death at 24 hours (D) or 48 hours (E) post-treatment.

### 14-3-3ζ inhibits BAD-induced cell death in CRC cells

To explore the potential of identified hits to treat diseases by triggering apoptosis in aberrant cells, we selected CRC as our disease model. We first determined whether the disruption of 14-3-3 protein:BAD PPIs could induce cell death in CRC cells. Two colorectal cell lines, Caco-2 and HT-29, were transfected with either BAD-mCitrine or BAD-AA-mCitrine, followed by incubation with FTY-720 or R18. BAD translocation to mitochondria following 14-3-3 protein inhibition was similar to what was observed in NIH-3T3 cells (**Figures 1; 5A, 5B**). Additionally, cell death was assessed following the over-expression of BAD or BAD-AA. After 72 hours post-transfection, a significantly higher degree of cell death was observed in cells co-expressing 14-3-3ζ and BAD-AA, compared to those only expressing 14-3-3ζ (control) or co-expressing 14-3-3ζ and BAD (**Figure 5C, 5D**). These observations imply that when 14-3-3ζ is unable to sequester BAD due to S112/136A mutations or due to the presence of 14-3-3 inhibitors in CRC cells, BAD translocates to the mitochondria to induce cell death.

**Figure 5.**
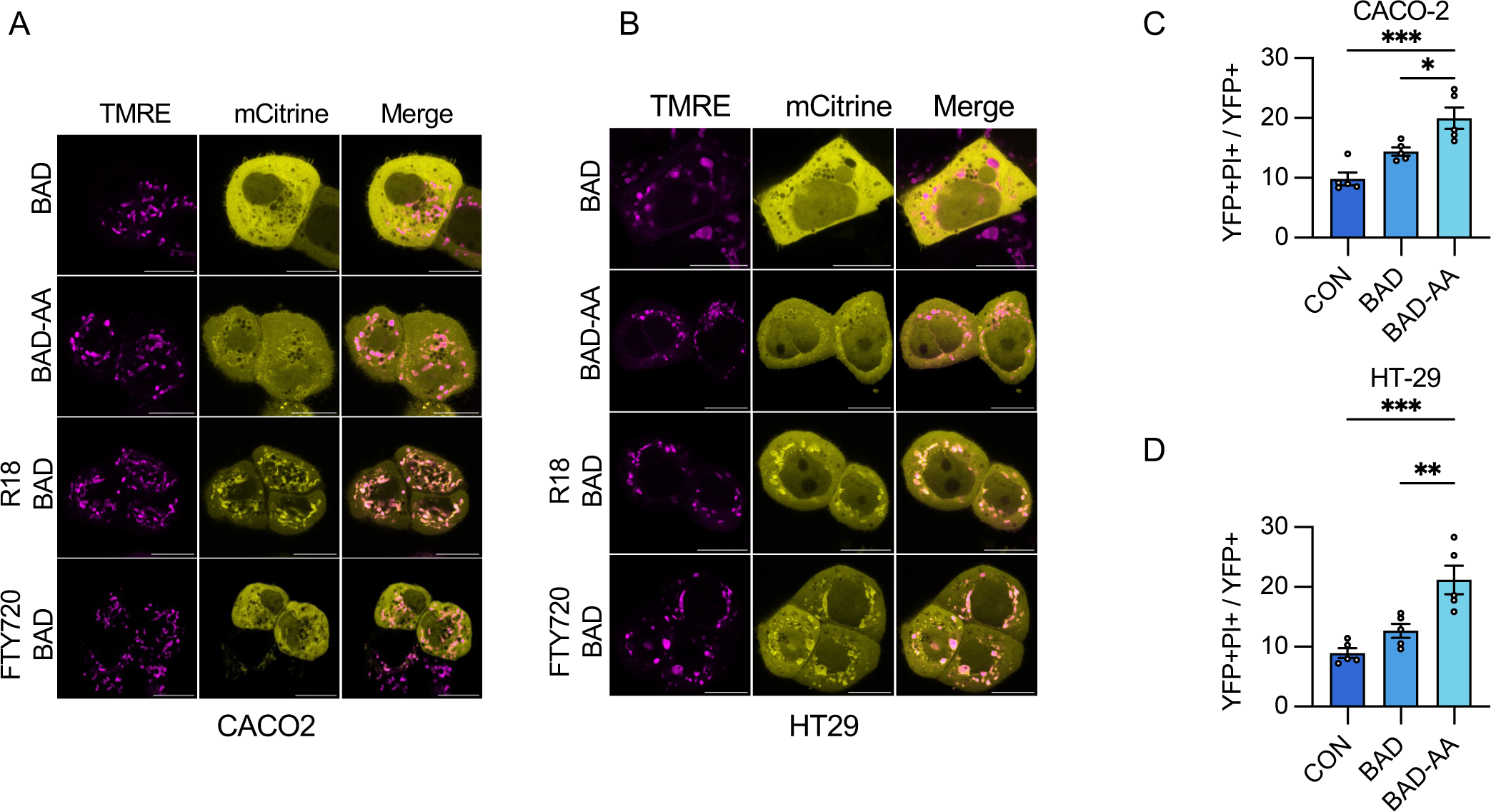
Over-expression of BAD leads to the death of colorectal cancel cells (CRCs) (**A**,**B**) Caco-2 (A) and HT-29 (B) were made to express mCitrine-conjugated BAD (BAD) or BAD mutants harboring S112A and S136A double mutations (BAD-AA). BAD-expressing cells were treated with either R18 (10uM) or FTY720 (2uM) for 24 hours. The mitochondrial membrane potential sensor TMRE (100 nM) was added to the medium 1 hour prior to live-cell imaging. Representative images were selected from three independent experiments, with at least five images were captured per dish in each experiment. (**C**,**D**) Caco-2 (C) and HT-29 (D) cells were transfected with either pBI-14-3-3ζ-Rluc8-mCitrine, pBI-14-3-3ζ-Rluc8-BAD-mCitrine, or pBI-14-3-3ζ-Rluc8-BAD-AA-mCitrine and allowed for 72 hours for plasmid expression. Cell death were evaluated using the Operetta CLS system. BAD-induced cell death was quantified as the ratio of YFP- and propidium iodide (PI)-double positive cells to YFP-positive cells. Data were collected from five independent experiments, each with triplicate measurements. (Statistical significance was determined using one-way ANOVA with a Dunnett’s post hoc test. \**P* < 0.05; \*\**P* < 0.01; \*\*\**P* < 0.001)

### Examination of hits in CRC cells

Among the 41 identified lead PADs from our primary screen in NIH-3T3 fibroblasts, some were immediately found to be unsuitable for systemic administration if re-purposed as chemotherapy. For example, crystal violet is a synthetic dye used for cell staining^[37]^; whereas some PADs are topical treatments, such as cetylpyridinium chloride, benzethonium chloride, and thonzonium bromide^[38–40]^. Various PADs are already used as chemotherapeutics, such as entrectinib, ceritinib, and ponatinib^[41]^. After excluding these PADs, 25 were assessed on Caco-2 and HT-29 CRC cells. Of these, 15 PADs caused more than 30% of cell death at 24 hours or 48 hours in both cell lines (**Figure 6A**), and further filtering of pro-drugs, salts, and potency narrowed our list to 13 PAD candidates (**Figure 6A**). Dose-response studies were then conducted with these 13 agents (**Figure 6B-6N**). Lomibuvir, terfenadine, penfluridol, and lomitapide were found to be the most effective, as they significantly induced cell death at concentrations as low as 5 µM (**Figure 6G, 6I, 6L, 6N**). Although lomibuvir consistently induced cell death at different concentrations, its efficacy in the magnitude of cell death attained was inferior to other candidates (**Figure 6N**), and it was excluded from further study.

**Figure 6.**
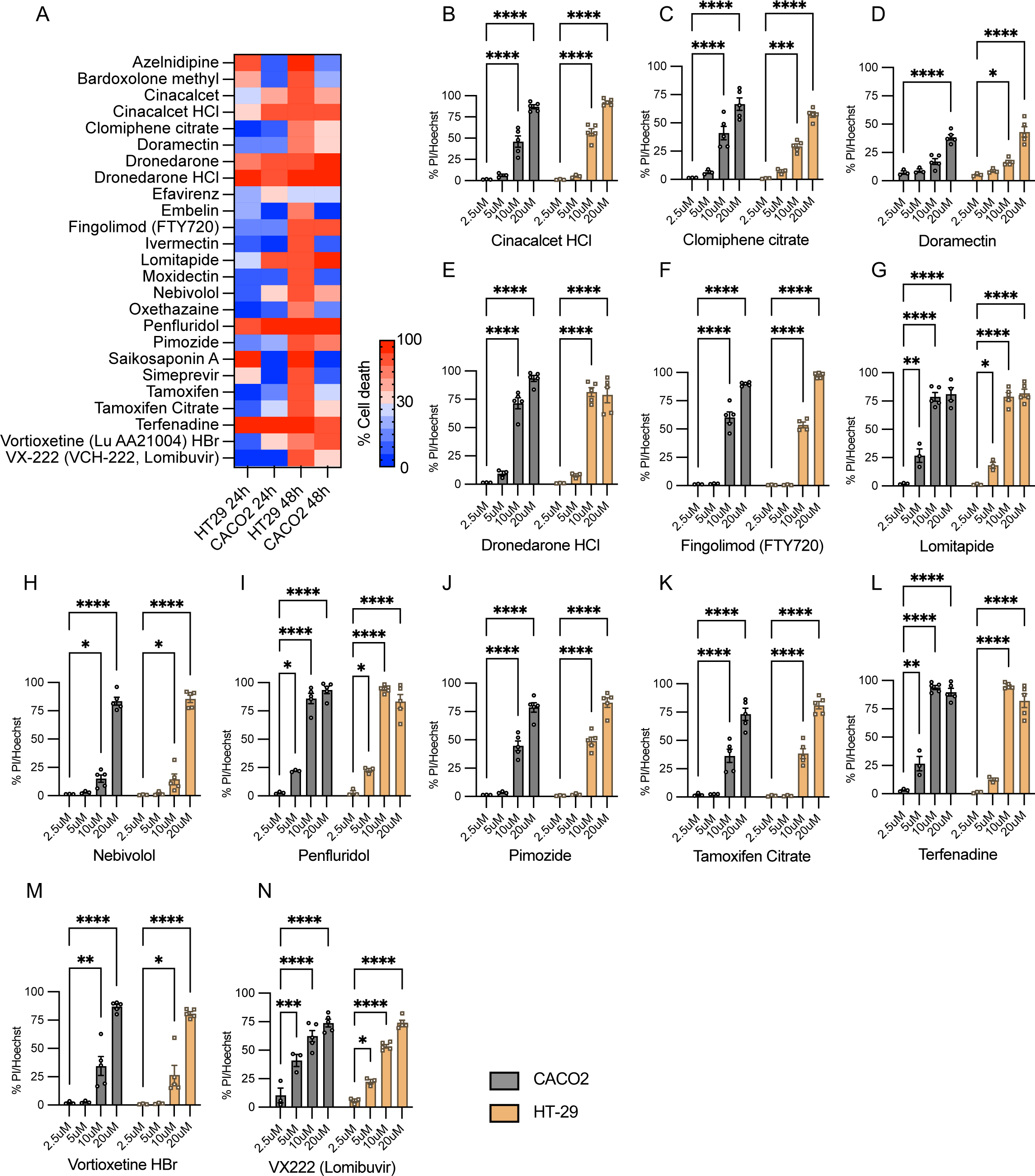
Capacity of identified hits to induce cell death in CRC cells. (A) Following the exclusion of unsuitable PADs, 25 hits were examined for their capacity to induce cell death In HT-29 and Caco-2 cells at 24 hours and 48 hours post-treatment, respectively. The heatmap shows that 15 of these hits induced more than 30% of cell death in both types of CRC cells. (**B-N**) After the filtering of pro-drugs, salts, and potency, does-response studies were performed for 13 selected hits. Lomitapide, terfenadine, penfluridol, and lomibuvir demonstrated effectiveness at concentrations as low as 5 µM. Data were collected from three (2.5 µM and 5 µM) or five (10 µM and 20 µM) independent experiments, each with duplicate measurements. (Statistical significance was determined using two-way ANOVA with a Tukey post hoc test. *#P* < 0.05 when compared with 2.5uM treatment; \**P* < 0.05; \*\**P* < 0.01; \*\*\**P* < 0.001; \*\*\*\**P <* 0.0001)

### Verifying the capacity of terfenadine, penfluridol, and lomitapide to induce apoptosis in cancer cells

Different pathways, such as apoptosis, necrosis, and autophagy, can account for cell death^[1]^. To ensure that these lead PADs induced apoptotic cell death via the disruption of 14-3-3 protein:BAD PPIs, we further assessed the relationship between apoptosis via hoechst/propidium iodide/annexin V incorporation and the mitochondrial translocation of BAD upon drug exposure. Given that caspase activation is a hallmark of apoptosis, a pan-caspase inhibitor, Z-VAD-FMK, was used to attenuate lead PAD-induced apoptosis. According to time courses of PI and Annexin V incorporation (**Figures 7A, B, F, G, K, L**), differences in the kinetics of cell death or apoptosis were seen with lead PADs in HT-29 and Caco-2 cells, such that terfenadine and penfluridol were able to induce cell death and apoptosis more rapidly than lomitapide. To assess caspase-dependent apoptosis, HT-29 and Caco-2 cells were treated for 24 hours with terfenadine (10uM) (**Figure 7C, 7D**) and penfluridol (10uM) (**Figure 7H, 7I**), or for 48 hours for lomitapide (10uM) (**Figure 7M, 7N**), in addition to the presence of Z-VAD-FMK or the control inhibitor Z-FA-FMK. Significantly diminished lead PAD-induced propidium iodide and/or annexin-V incorporation was observed in cells pre-treated with Z-VAD-FMK compared to those pre-treated with either DMSO or a control inhibitor Z-FA-FMK, implying caspase activation following lead administration. Additionally, confocal imaging showed that mitochondrial translocation of BAD occurred after the addition of lead PADs (**Figure 7E, 7G, 7O**), demonstrating that disruption of 14-3-3:BAD PPIs by lead PADs induces apoptosis.

**Figure 7.**
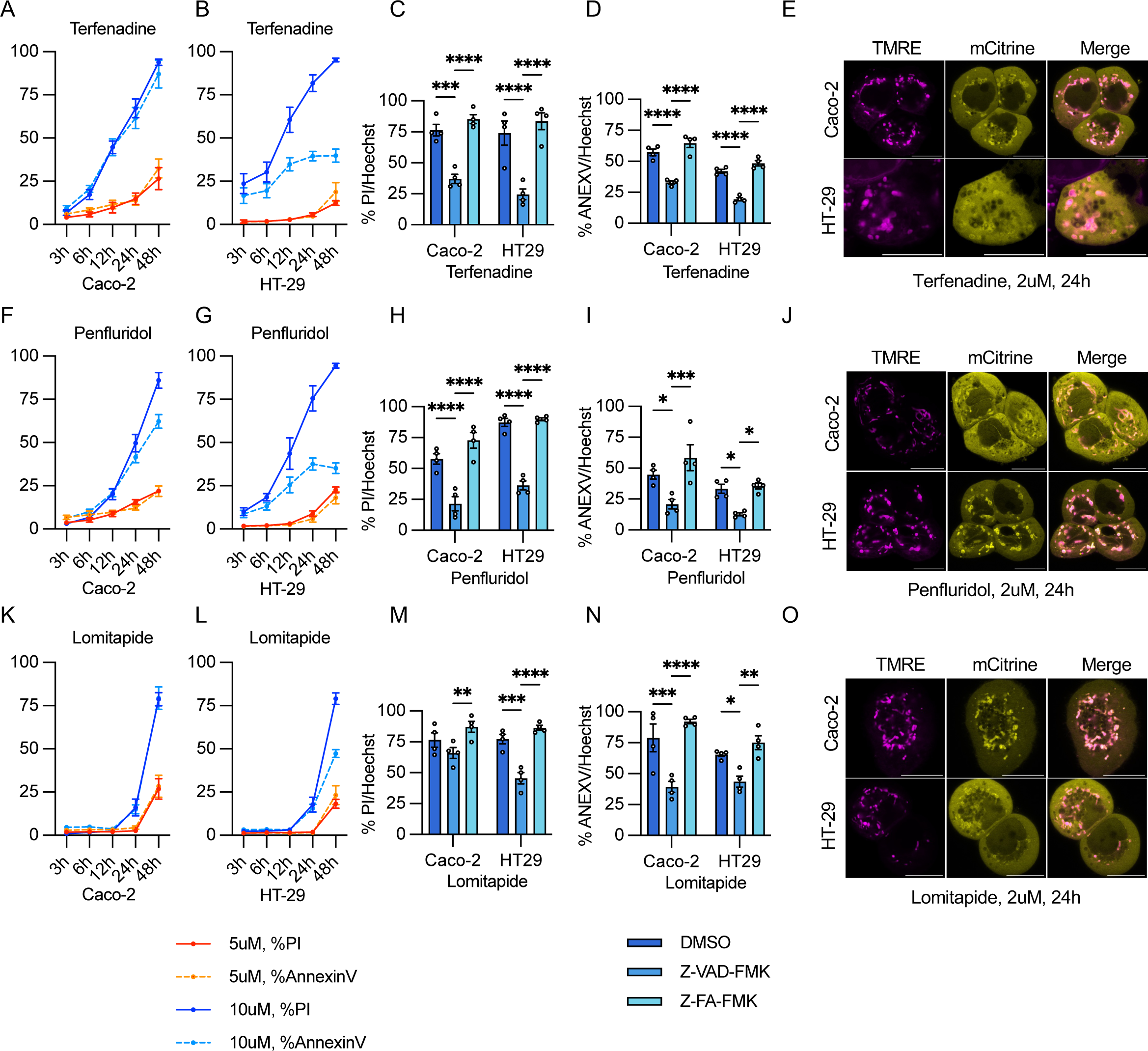
Ability of lead hits to induce apoptotic cell death in colorectal cancer cells. (**A, B, F, G, K, L**) Time-response studies using Hoechst/propidium propidium iodide (PI)/annexin V (ANEXV) incorporation assays were conducted for terfenadine (A, B), penfluridol (F, G), and lomitapide (K, L) in Caco-2 (A, F, K) and HT-29 (B, G, L) cells at concentrations of 5 µM and 10 µMData were collected from three (5uM) or five (10uM) independent experiments, each with duplicate measurements. (**C, D, H, I, M, N**) The pan-caspase inhibitor Z-VAD-FMK or its control inhibitor Z-FA-FMK (both at 20 µM) was introduced to determine the mechanism of cell death induced by lead PADs. PI incorporation (C, H, M) and annexin V incorporation (D, I, N) were measured in Caco-2 or HT-29 cells after 24 hours (C, D, H, I) or 48 hours (M, N) of treatment. Data were collected from four independent experiments, each with duplicate measurements. Statistical significance was determined using two-way ANOVA with a Tukey post hoc test. \**P* < 0.05; \*\**P* < 0.01; \*\*\**P* < 0.001; \*\*\*\**P <* 0.0001. (**E**, **J**, **O**) Mitochondrial translocation of BAD was obeserved in CRC cells treated with terfenadine (E), penfluridol (J), and Lomitapide (O) at 2 µM, 24 hours post-treatment. Representative images were selected from three independent experiments, with three images were captured per dish in each experiment. (Scale bar= 10 µm)

## Discussion

Cancer arises from uncontrolled cell proliferation, and one of the primary goals of chemotherapies is to induce cancer cell death. The overexpression of 14-3-3ζ and its related isoforms in the context of cancer has long been associated with poor clinical outcomes due to increased cell survival ^[15, 42]^. Although it has been demonstrated that inhibiting 14-3-3ζ triggers apoptosis in cancer cells, there are currently no approved chemotherapies that target 14-3-3ζ:BAD interactions^[17]^. The primary aim of this study was to explore the possibility of identifying anti-CRC compounds by their capacity to interrupt 14-3-3ζ:BAD interactions, and this was executed by the generation of a biosensor capable of detecting 14-3-3ζ:BAD PPIs. While a cell-free compound screen has been used in discovering inhibitors of 14-3-3ζ:BAD PPIs, limited data exist on whether these drugs are able to induce apoptosis in living cells or animal models^[24]^. To address this limitation, we developed a BRET-based biosensor that was capable of detecting 14-3-3ζ:BAD PPIs in living cells.

In this study, we utilized a short fragment of murine BAD to construct the BRET sensor rather than the full-length BAD. Previous research has demonstrated that BAD overexpression leads to apoptosis in various cell types^[43, 44]^, and an advantage of using the BAD-112-136F is its inability to interact with its client BCL-2 proteins due to the lack of BH-3 domain, leading to reduced cytotoxicity^[45]^. Furthermore, it was not possible to detect BRET when the Rluc8 acceptor and mCitrine donor were fused to 14-3-3ζ and full-length BAD, respectively. Since energy transfer between Rluc8 and mCitrine requires a distance shorter than 10nm, we assumed that the inability to detect BRET was due to the distance between the fusion sites and interaction sites. Indeed, in another study using time-resolved fluorescence resonance energy transfer (FRET), a technique where a fluorescent protein is used as an energy donor instead of luciferase, the FRET acceptor was directly fused to the serine residue that mediates interactions^[24]^. Therefore, different truncated forms of BAD were generated to identify the optimal fusion strategy for measuring BRET efficiency between 14-3-3ζ-Rluc and BAD-truncate-mCitrine. Given that the interactions between 14-3-3ζ and BAD occur between the C-terminus of 14-3-3 and S112 or/and S136 of BAD, it was not surprising that the combination of 14-3-3ζ-Rluc8 and 112-136F-mCitrine represented the optimal combination in the constructing the BRET sensor^[12, 46]^.

Since it was uncertain if the BAD-112-136F could represent the full-length BAD in its interactions with 14-3-3ζ, further evaluations were conducted by introducing mutations at Ser-Ala mutations at S112 and/or S136. Unlike the S112A mutation, S136A significantly disrupted the association between 14-3-3 and BAD-112-136F, as indicated by reduced BRET. This aligns with a prior study suggesting that S136, rather than S112, primarily mediates 14-3-3:BAD interactions^[47]^. Additionally, in contrast to 112-136F-AA, 112-136F is specifically sequestered in the cytoplasm. This indicates that 14-3-3ζ interacts with this truncated form similarly to how it interacts with full-length BAD, but only if the serine residues crucial for 14-3-3:BAD interactions remain intact.

To assess the capacity of this sensor to discover disruptors of 14-3-3(:BAD PPIs, we introduced two well-known 14-3-3 inhibitors, FTY720 and I-2,5, and both compounds signficantly reduced BRET. It is worth mentioning that we did not assess if there were any differences in affinity between 14-3-3(:BAD and 14-3-3(:112-136F. Nevertheless, drugs identified to disrupt 14-3-3(:112-136F PPI should be effective, as this smaller fragment likely accesses the binding groove of 14-3-3ζ more readily than the full-length BAD^[48]^. Additionally, it is worth mentioning that the high homology among different 14-3-3 isoforms permits them to share client proteins and form homo- or hetero-dimers, suggesting that the identified PADs are highly likely to disrupt PPIs not only between BAD and 14-3-3 (but also between BAD and other 14-3-3 isoforms^[46]^. An important caveat of our reporter system is that we cannot distinguish if PADs directly block or disrupt the amphipathic groove of 14-3-3 (where PPIs occur or if PADs promote the dephosphorylation of Ser112 and Ser136F on the BAD fragment^[10] [43, 44]^. Additional studies are required to examine the mechanisms of action of each lead PAD.

The efficiency of utilizing HTS to develop novel anti-cancer compounds has been underscored by the discovery of sorafenib, palbociclib, and ABT-199^[49–52]^. However, a recognized drawback of this drug discovery approach is the increased risk of false positives and false negatives due to the lack of replication and the use of miniaturized reaction systems^[53, 54]^. To increase the chances of identifying potential compounds, we first ensured the robustness of our sensor by achieving a Z-factor greater than 0.5^[30]^. Secondly, to minimize false negatives, we tested each compound twice at four different concentrations in our primary screens but only recorded the highest BRET reduction for each concentration.

After evaluating the capacity of identified compounds to induce cell death in NIH-3T3 fibroblasts, a group of drugs that decreased BRET by more than 34% and triggered more than 30% of cell death emerged as potential hits capable of killing target cells by disrupting 14-3-3(:BAD PPIs. Interestingly, this BRET reduction aligns with that caused by 112-136F-AA-mCitirine, suggesting that the ability of a compound to completely dissociate the 14-3-3(:112-136F complex is indicative of its potential to induce cell death. Another group of drugs also arose from our screens such that they were capable of reducing BRET reduction by more than 34%, but without notable efficacy in inducing cell death in NIH-3T3 cells. A possible explanation for this is that our screening is based on a cell-based assay^[36]^, and the complex intracellular environment makes it challenging to determine whether the dissociation of 14-3-3(:BAD resulted from a direct inhibitory action on 14-3-3 (or BAD, or an indirect effect on the upstream signaling pathways that promote 14-3-3(:BAD interactions. Therefore, other than disrupting 14-3-3:BAD interactions, these compounds may have additional effects, such as up-regulating anti-apoptotic BCL-2 proteins, which promote cell survival^[11]^.

Although the altered expression of 14-3-3 in CRC has been reported in several studies, the role of 14-3-3(:BAD in the survival of CRC cells remains unclear^[17, 55, 56]^. Our research provides the first evidence that disruption of 14-3-3(:BAD PPIs can promote CRC cell death. We used two representative CRC cell lines, Caco-2 and HT-29, to validate our assay^[57–59]^. Terfenadine, penfluridol, and lomitapide were selected as lead compounds after conducting dose-response studies with 13 selected compounds. Additionally, we also collected data on the efficacy of all 101 compounds that were identified from the primary screen in inducing cell death in these two CRC cell types at 20uM (**Supplementary table 1**). These data would be invaluable for future research exploring the different capacities of drugs to target CRC cells.

To confirm that lead PADs induce apoptotic cell death by disrupting 14-3-3(:BAD PPIs, we further tested whether inhibiting caspase activation could mitigate lead PAD-induced cell death and if these lead PADs could promote the mitochondrial translocation of BAD. In most cases, Z-VAD-FMK treatment prevented increases in PI-positive and Annexin-V-positive cells; however, in lomitapide-treated Caco-2 cells, Z-VAD-FMK had no effect on propidium iodide incorporation, despite preventing annexin V incorporation (**Figure 7M**). This is likely due to the kinetics of lomitapide in Caco-2 cells whereby the early stages of apoptosis, marked by the binding of annexin V to phosphatidylserine, is being observed, without a loss of membrane integrity that is needed for propidium iodide entry into the cell^[60]^. As Z-VAD-FMK cannot inhibit necroptosis, a process where cells shift to necrosis when they cannot complete apoptosis^[61, 62]^, it is possible that other forms of cell death may also be occuring. Nevertheless, confocal imaging showed that all three lead PADs trigger the mitochondria translocation of BAD, confirming apoptotic cell death.

With the variability of 14-3-3 protein expression across individuals, a personalized medicine approach could be undertaken to explore the potential of lead PADs to treat CRC by careful evaluation of tissues from people living with CRC^[56]^. A significant advantage of our screening strategy is our focus on repurposing drugs from PADs. Therefore, all of our identified hits have been tested for their safety in prior phase 1 clinical trials. Interestingly, penfluridol and lomitapide have been previously suggested to have potential for treating CRC, but their mechanisms were not fully defined^[63–65]^. Our study not only further validates their therapeutic value but also provides insight into the mechanism by which these drugs may ameliorate CRC. Nevertheless, additional in-depth pre-clinical studies in animal models are required. We recognize that PADs may also have effects in other cell types, and improving cell type specificity is clearly warranted^[8]^. A potential approach could also be to adopt a localized use of PADs to treat colorectal tumors, wherebyregional drug administration might also enhance the specificity of these compounds^[66]^.

## Conclusion

With the use of a novel BRET-based sensor to monitor 14-3-3(:BAD interactions in living cells, we successfully identified terfenadine, penfluridol, and lomitapide as having the abilities to disrupt 14-3-3(:BAD interactions and induce apoptosis of CRC cells. Although further research is critical to validate the ability of these compounds to ameliorate CRC in animal models and in humans, these hits represent potential chemical backbones that can be modified and translated into new chemical entities for the treatment of CRC. In addition, our screening approach has shown significant potential for the discovery of novel therapeutics for the treatment of other apoptosis-related diseases.

## Acknowledgments

The authors would like to thank Drs. Jace Jones-Tabah and Terry Hébert (McGill University) for providing the protocol and original Rluc8 vectors for the generation of the BRET-based sensor. We would also like to thank Dr. Aurélie Cleret-Buhot of the Cell Imaging core facility of the CRCHUM for performing the confocal microscopy acquisitions, Dr. Alexis Vivoli (CRCHUM) for data visualization assistance, George Vornicu (University of Montreal) for conducting preliminary cell death assays, and Dr. Petronela Ancuta (CRCHUM) for providing the Caco-2 and HT-29 cell lines. The authors also thank the Cellular physiology core facility (CRCHUM) for their help with the Operetta CLS and the Cellular imaging core facility (CRCHUM) for their help with confocal imaging.

## Supplementary information

1. Supplementary figure 1-Truncated forms of BAD used in the development of the BRET sensor

---

**Supplementary Figure 1.**
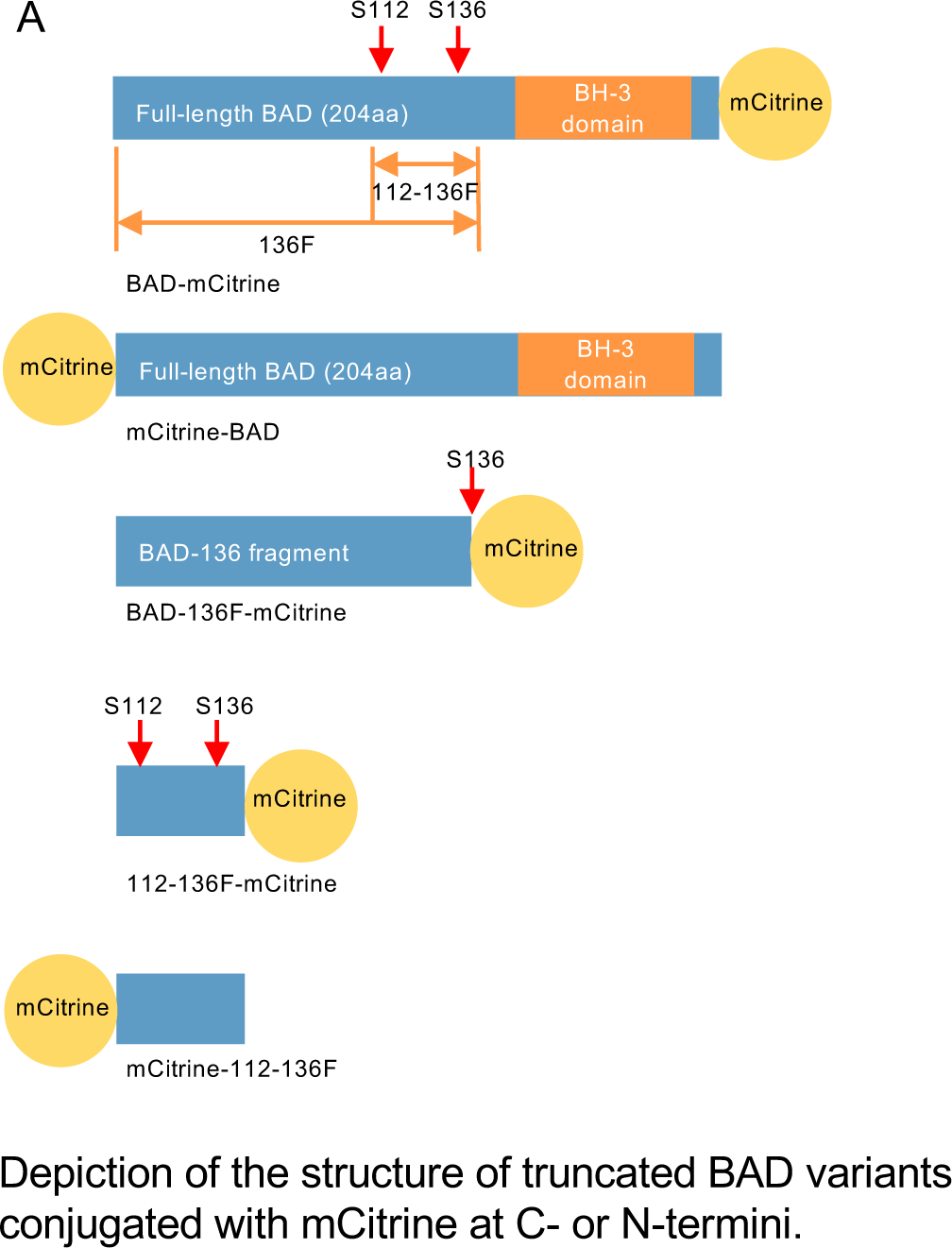
Truncated forms of BAD.

